# Cytogenetically-based biodosimetry after high doses of radiation

**DOI:** 10.1101/647362

**Authors:** M. Pujol-Canadell, J.R. Perrier, L. Cunha, I. Shuryak, A. Harken, G. Garty, D.J. Brenner

## Abstract

Dosimetry is an important tool for triage and treatment planning following any radiation exposure accident, and biological dosimetry, which estimates exposure dose using a biological parameter, is a practical means of determining the specific dose an individual receives. The cytokinesis-blocked micronucleus assay (CBMN) is an established biodosimetric tool to measure chromosomal damage in mitogen-stimulated human lymphocytes. The CBMN method is especially valuable for biodosimetry in triage situations thanks to simplicity in scoring and adaptability to high-throughput automated sample processing systems. While this technique produces dose-response data which fit very well to a linear-quadratic model for exposures to low linear energy transfer (LET) radiation and for doses up for 5 Gy, limitations to the accuracy of this method arise at larger doses. Resolution at higher doses is limited by the number of cells reaching mitosis. Whereas it would be expected that the yield of micronuclei increases with the dose, in many experiments it has been shown to actually decrease when normalized over the total number of cells. This variation from a monotonically increasing dose response poses a limitation for retrospective dose reconstruction. In this study we modified the standard CBMN assay to increase its resolution following exposures to higher doses of photons or a mixed neutron–photon beam. The assay is modified either through inhibitions of the G2/M and spindle checkpoints with the addition of caffeine and/or ZM447439 (an Aurora kinase inhibitor), respectively to the blood cultures at select times during the assay. Our results showed that caffeine addition improved assay performance for photon up to 10 Gy. This was achieved by extending the assay time from the typical 70 h to just 74 h. Compared to micronuclei yields without inhibitors, addition of caffeine and ZM447439 resulted in improved accuracy in the detection of micronuclei yields up to 10 Gy from photons and 4 Gy of mixed neutrons-photons. When the dose-effect curves were fitted to take into account the turnover phenomenon observed at higher doses, best fitting was achieved when the combination of both inhibitors was used. These techniques permit reliable dose reconstruction after high doses of radiation with a method that can be adapted to high-throughput automated sample processing systems.

## INTRODUCTION

In the event of large-scale radiation exposure due to an improvised nuclear device (IND), biological dosimetry is an important tool to determine the dose received by a given individual. One method of biological dosimetry is the scoring of radiation-induced chromosomal damage in peripheral blood lymphocytes, known as the dicentrics assay, which can provide a reliable and independent assessment of dose (1). Although this assay is considered the gold standard in biodosimetry, dicentrics scoring usually requires analysis by technically skilled personnel and is therefore not quick enough for time-sensitive triage situations. These triage situations, which could occur following radiation exposure emergencies, would need to process a large number of victims and produce dosimetric data for medical management purposes.

A promising technique for biological dosimetry in triage situations is the *in vitro* cytokinesis-blocked (CB) micronucleus (Mn) assay, originally developed by Fenech and Morley (2). The CBMN assay quantifies the frequency of micronuclei (Mn) in binucleated cells (BNCs) derived from peripheral lymphocytes. Whole chromosomes, as well as chromosome fragments, can be in micronuclei that lag behind at anaphase during nuclear division; the Mn form into small, rounded bodies surrounded by their own nuclear envelope and can persist up to 1 year after irradiation (3). The CBMN is a particularly useful biodosimetric tool for quantifying radiation-induced chromosomal damage for population triage due to the simplicity of Mn scoring and the availability of high-throughput fully automated systems (4)(5)(6)(7)(8). However, The accuracy of cytogenetically-based techniques such as CBMN for dose reconstruction decreases for doses higher than 5 Gy of purely photon irradiations (9)(10)(11). At doses > 5 Gy, many of the highly damaged cells experience delays in the cell cycle (12) or are unable to reach mitosis, which reduces the number of scorable binucleated cells, and hence the overall number of Mn. The Mn frequency increases monotonically up to about 5 Gy, but at higher doses it does not increase as expected but rather decreases with increasing dose. The result at these high doses is a “turnover” in the dose response curve, which will lead to inaccuracies in dose estimation with an under-representation of the absorbed dose.

High dose biodosimetry has been a persistent problem, specifically for the CBMN assay, and few studies have attempted to address this problem. Müller and Rode (10) assessed up to 15 Gy using CBMN by focusing on the analysis of Mn per micronucleated BN cells, and on the ratio of trinucleated to tetranucleated cells, but their analysis was limited since cells were only tested from a single donor. A study from Kacprazak et al. (9) also assessed Mn per micronucleated BN cells, and on the ratio of trinucleated to tetranucleated cells, from three different donors and concluded that these parameters exhibit large variations between individuals. Large interindividual differences at high doses were also found when the ratio of Mononucleated/Binucleated was considered (11). These two studies suggest that to avoid dose underestimations, proliferation assays should rather be used at relatively high doses. However, proliferation tests depend on culture conditions as well as differences between individuals, reducing their utility for dose reconstruction.

Previous studies have shown that cell cycle arrest induced by high radiation doses can be significantly reduced by treatment with caffeine, which abrogates the G2/M checkpoint allowing cell cycle progression into metaphase (13)(14)(15)(16)(17)(18)(19)(20). This treatment has been applied for dicentric analysis in biodosimetry purposes after high doses of low LET irradiations (21)(18)(19)(20). With this study we aim to apply this same approach of cell cycle arrest to the CBMN assay, in an effort to extend the monotonic section of the dose response and therefore the utility of this assay. Inhibition of the G2/M and spindle checkpoints was performed using caffeine and ZM447439 (an Aurora kinase inhibitor), respectively. The effects of these inhibitions were tested in samples exposed to up to 10 Gy of photons and up to 4 Gy of neutrons. Overall, these studies show that with simple modifications to the standard CBMN protocol we are enable dose reconstruction at high doses thereby extending the applicability of this cytogenetically-based assay to many radiation accident scenarios where it is likely that individuals will be exposed to higher doses.

## MATERIALS AND METHODS

### Blood Collection and Irradiation

Peripheral blood from 8 (4 female and 4 male) apparently healthy volunteers, aged between 29 and 48 y, with no history of exposure to ionizing radiation or clastogenic agents, were obtained after they signed an informed consent. The study was reviewed by Columbia University’s Institutional Review Board (IRB) and the use of human materials was approved under the IRB protocol AAAF2671. Blood was collected by venipuncture into vacutainer tubes containing sodium heparin (Becton Dickinson, Franklin Lakes, NJ). The total volume of blood collected from each donor was 20 ml.

For photon irradiation, samples of whole blood were individually irradiated using a Gammacell 40 ^137^Cs irradiator (Atomic Energy of Canada, Ltd., Chalk River, Canada) at a dose rate of 0.73 Gy/min. The ^137^Cs irradiator dose rate is verified annually with thermoluminescent dosimeters (TLDs). The homogeneity of the exposure across the sample volume is verified using EBT3 Gafchromic™ film (Ashland Advanced Materials, Bridgewater, NJ), which indicated less than 5 % variation within the sample. Samples were exposed to 2, 4, 6, 8 and 10 Gy. Control samples were sham irradiated and received zero dose.

Neutron irradiations were performed at the Columbia IND Neutron Facility (CINF)(22). This irradiator is capable of exposing cells, blood, or small animals to neutrons with an energy spectrum mirroring that produced at 1 to 1.5 km from the epicenter of the Hiroshima detonation (23), a distance where many exposed survivors would be expected in the event of an IND detonation. In our initial biodosimetry studies, we found these neutrons to have a relative biological effectiveness (RBE) of about 4 for induction of micronuclei (22). Further details of the neutron spectrum (23) and dosimetry (22) have been previously published.

For neutron irradiations, 3 ml vacutainers were placed in custom-made sample holders during exposure (24). 16 tubes were loaded on an 18-position Ferris wheel, centered on the beam axis, with two water containing tubes loaded at the empty positions. The Ferris wheel is positioned such that neutrons, emitted from a thick beryllium target at 60° to the beam axis, hit the center of each tubes. During irradiation, the tubes are rotated around the beam axis at 0.5 RPM, to average out 20% angular variations in dose rate, and flipped end to end half way through the exposure, to ensure an equal dose to the whole sample.

To generate neutrons, 20 μA of a mixed Hydrogen/deuterium beam impinged on a thick beryllium target generating a dose rate of 2.2 Gy/h neutrons with a concomitant γ dose of 0.5 Gy/h (18%) at the position of the sample tubes. For simplicity we will refer to the resulting radiation beam as “neutrons”.

Four tubes (1 per donor) were irradiated with 0, 1, 2, 3, or 4 Gy delivered in 1 Gy fractions; the beam was stopped for a few minutes between fractions to flip tubes and remove those that had reached the prescribed dose. Whenever tubes were removed from the wheel, two tubes containing water blanks were placed at either end of the string of blood samples in order to ensure a uniform scatter dose to the irradiated tubes.

### Lymphocyte Culture and Slide Preparation

Whole blood samples (300 uL) were diluted in 2.7 mL of PB-MAX™ Karyotyping media (Life Technologies, Grand Island, NY). Blood cultures were set up in a 48 well plate and cultured at 37°C in a humidified atmosphere with 5% CO_2_ for 44 h. Culture media was then exchanged with media containing cytochalasin B (Cyt-B; Sigma-Aldrich, St. Louis, MO) at a final concentration 6 µg/ml. The cells were cultured for an additional 26 hours to arrest cytokinesis and induce the formation of first-divided BN cells. For the caffeine treatments, caffeine (4 mM; Sigma-Aldrich, St. Louis, MO) was added at the same time as Cyt-B to abrogate the G2/M checkpoint. For the spindle checkpoint inhibition, ZM447439 (5 uM; Millipore, Burlington, MA) was added at 57 hours to abrogate the spindle checkpoint. This time point was chosen based on a previous work (18) that determined that it takes 57 hrs for 10-Gy irradiated lymphocytes to reach metaphase.

Cells were harvested at 70 hrs for the standard Mn assay and up to 78 for caffeine protocol optimization. For the dose response curves of photons and neutrons, cultures were harvested at 74 hrs after irradiation. The inhibitors were added to the culture separately or combined. To harvest the cultures, samples were incubated in hypotonic solution (KCl 0.075 M0.075 M KCl; Sigma-Aldrich) for 10 minutes, and fixed with methanol:glacial acetic acid, (5:1, v/v) (FisherThermo Fisher Scientific, Rockford, IL). Cells were dropped onto slides, allowed to air-dry, and finally stained with DAPI Vectashield® mounting medium (#H-1200; Vector Laboratories, Inc., Burlingame, CA).

### Microscope analysis

Entire slides were scanned at 10× magnification with Metafer 4 software (MetaSystems, Altlussheim, Germany) using a microscope (Carl Zeiss Microscopy GmbH, Jena, Germany) and motorized stage capable of scanning eight slides. The Metafer classifier was run on the Metafer MSearch platform version 3.5 software. Images were captured using a high-resolution, monochrome megapixel charge coupled device (CCD) camera.

### Statistical analysis

Statistical comparisons were performed between micronucleated/binucleated (MN/BN) values from different treatments using paired Student’s t tests, with p values < 0.05 considered significant. Dose response curve fitting was performed using nonlinear least squares (using the *nls* function in *R* 3.5.1) separately for each condition, weighted by the number of analyzed cells for each data point. The dose response curve was fitted using the following equation, based on the standard linear-quadratic model (1):

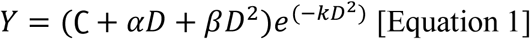

Here Y is the Mn/BN yield, C the background value, α and β are the linear and quadratic dose response parameters respectively, D is the dose, and an additional parameter k is included to represent the observed turnover at high doses, which is probably caused by mitotic arrest of heavily radiation-damaged cells. Goodness-of-fit was assessed by the coefficient of determination (R^2^) and root mean squared error (RSME).

## RESULTS

### Extended caffeine cultures optimization

Figure 1 shows yields of Mn/BN at different culture times (70, 74, and 78 h) compared to the standard CBMN protocol of 70 hrs after 0, 2, 4, 8 and 10 Gy of gamma radiation. The results show a significant higher frequency of Mn/BN at doses of 8 or 10 Gy for 74 and 78 hrs caffeine-treated cultures p<0.05, compared to the untreated cultures. The Mn/BN ratio at 78 h with caffeine were not significantly different than that measured at 74 hr assay with caffeine at any dose.

**Figure 1.**
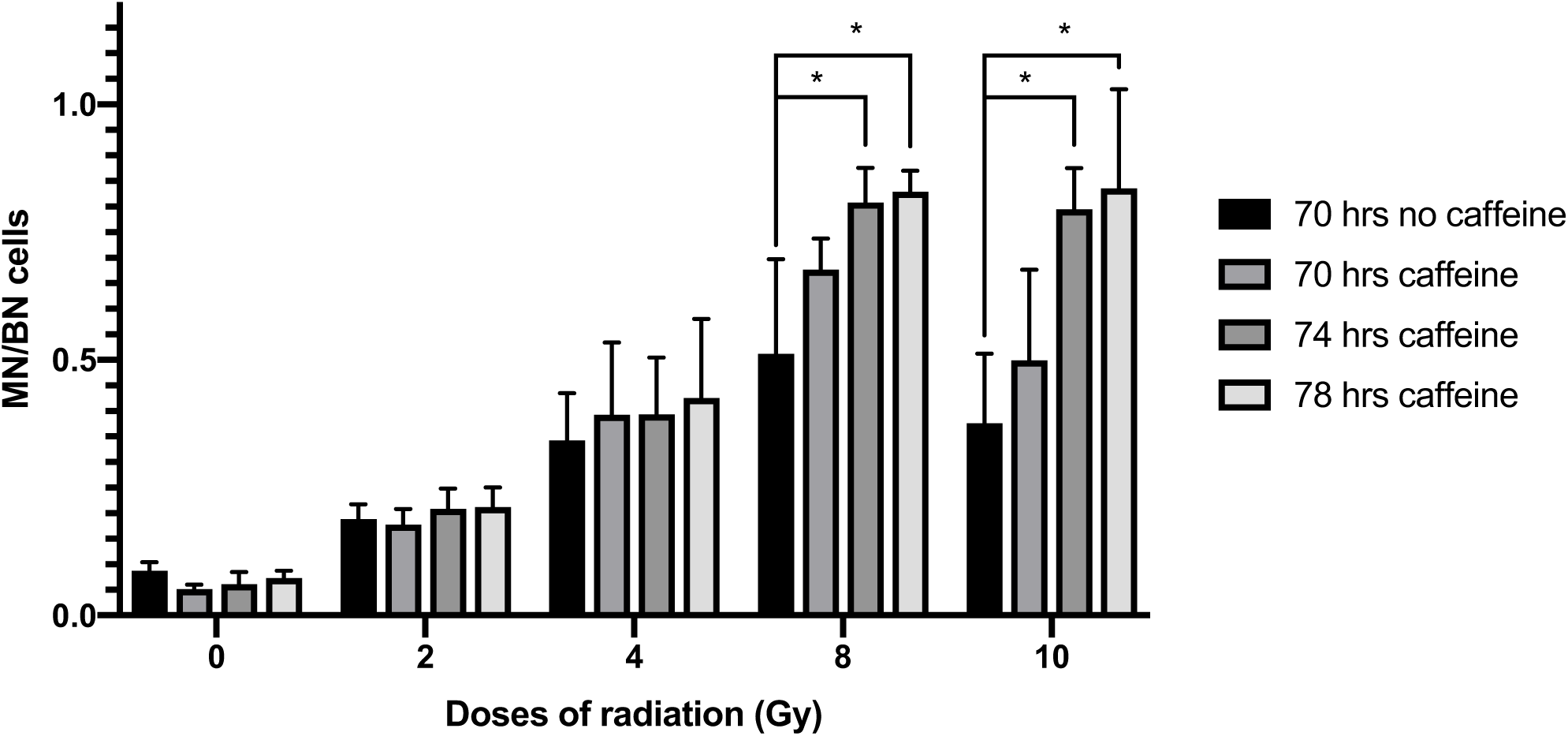
Micronuclei frequencies per binucleated cells after Caffeine culture optimization at different doses of radiation. Error bars show SD from the 4 different donors. * indicates significant differences p<0.05

### Effect of caffeine and ZM447439

Figure 2 shows the Mn/BN ratios after photon exposure (0 to 10 Gy; Figure 2A) and exposure to neutrons (0 to 4 Gy; Figure 2B). For both radiation types, we compared the standard 70 h CBMN protocol with 1) the 74 h caffeine-CBMN assay, 2) the 74 h ZM447439 and 3) the 74 h CBMN assay with the combination of caffeine and ZM447439. The results show that there was no significant difference in Mn frequency within assay conditions within assay conditions of 0 to 6 Gy of photons or for 0 to 1 Gy of neutrons. As the photon dose increased to 10 Gy, the combined effect of caffeine and the ZM447439 showed the largest yield of Mn/BN, although an increase was also observed when only caffeine or ZM447439 was added to the cultures (Fig. 2A). For the 3 Gy and 4 Gy neutron dose points (Fig. 2B), any of the three caffeine/ZM447439 treatments significantly (p < 0.05) improved the detection of Mn/BN, as opposed to the standard CBMN protocol that lead instead to a reduced Mn/BN ratio with increasing dose (black bars in Fig. 2B).

**Figure 2.**
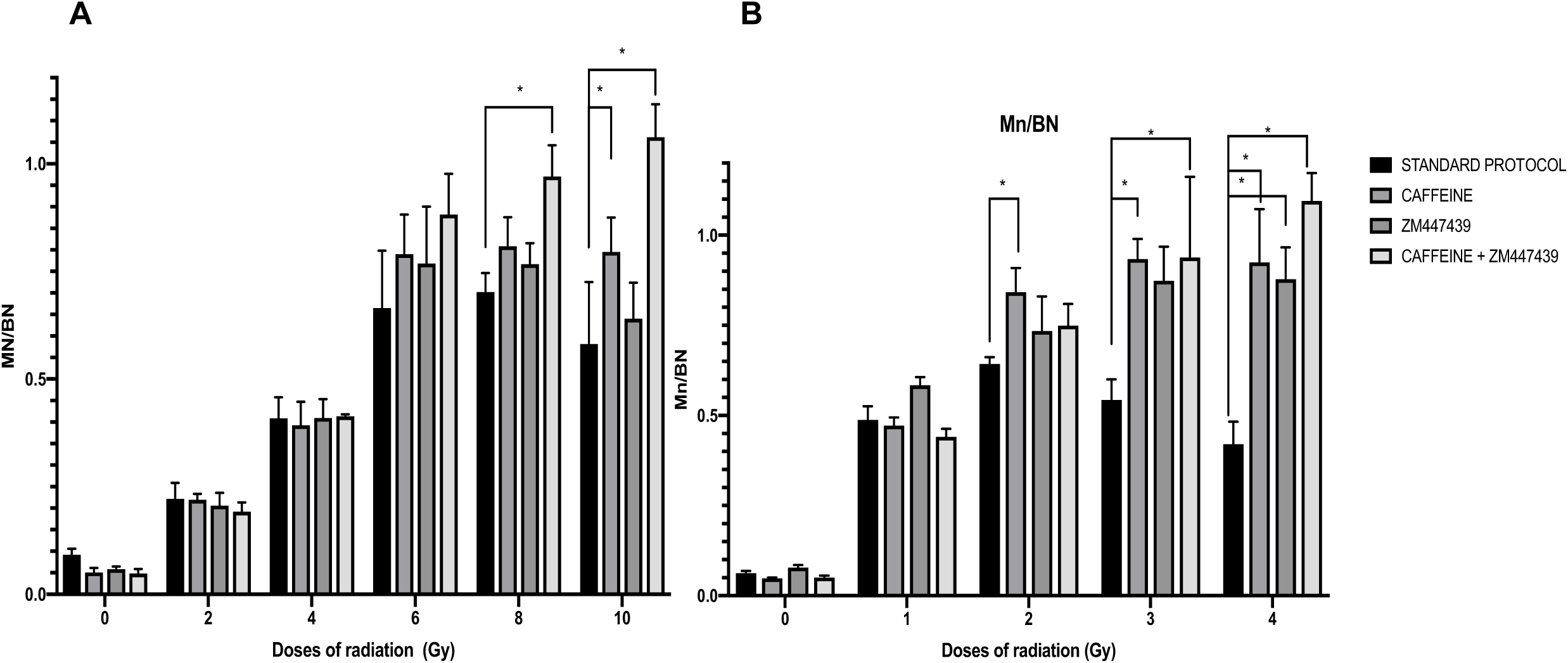
Micronuclei frequencies per binucleated cells. A (photon irradiation), B (neutron irradiation) at different doses of radiation. With four different treatments. Error bars show SD from the 4 different donors. * indicates significant differences p<0.05

### Dose effect curves

The data for the curve fitting using Equation 1 is shown in Table 1. Figures 3 and 4 show the dose response curves (n= 4 individuals) across the different conditions for photons and neutrons, respectively. As shown in Figures 3A corresponding to the standard CBMN protocol, the response curve starts to decrease at doses > 8 Gy. This turnover effect is more apparent after neutron exposures (Fig 4 A) where the Mn/BN ratio peaks at 2 Gy but decreases at higher doses. Overall, the dose response curves improved, indicated by a reduction of the k ‘‘turnover’’ parameter (Table 1), by optimizing the standard CBMN protocol with the addition of either caffeine and/or ZM447439 to the blood culture. One exception is in the photons irradiated samples treated only with ZM447439 where the k value is slightly higher than in the standard protocol. Condition D, which corresponds to the double inhibition of G2/M and spindle checkpoints showed the weakest turnover at high doses for both, photon and neutron irradiated samples (Fig. 3D and 4D).

**Figure 3.**
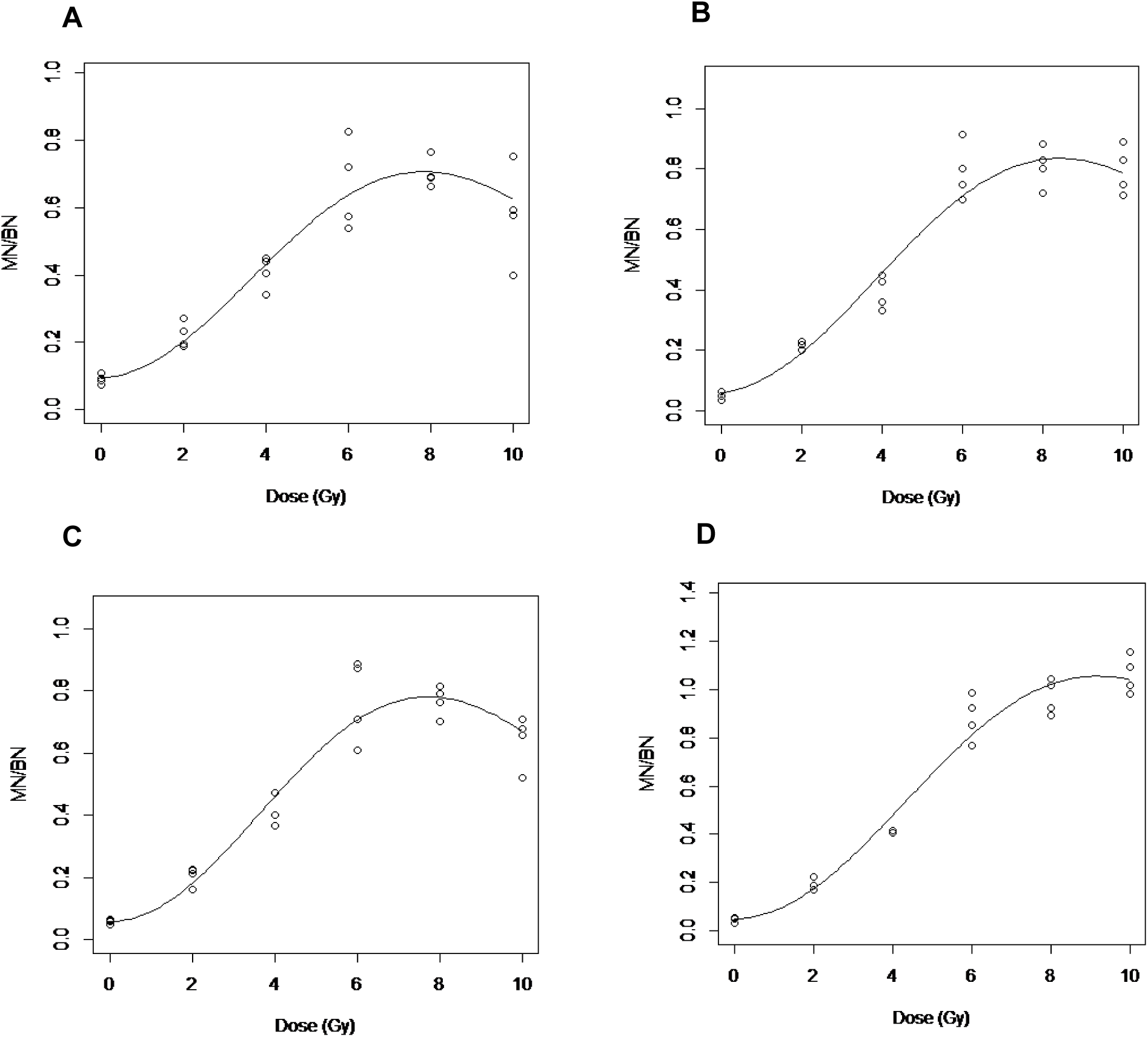
Dose effect curves after photon irradiations. A. Standard CBMN assay; B. Caffeine CBMN assay; C. ZM447439 CBMN; D. Caffeine & ZM447439 CBMN assay.

**Figure 4.**
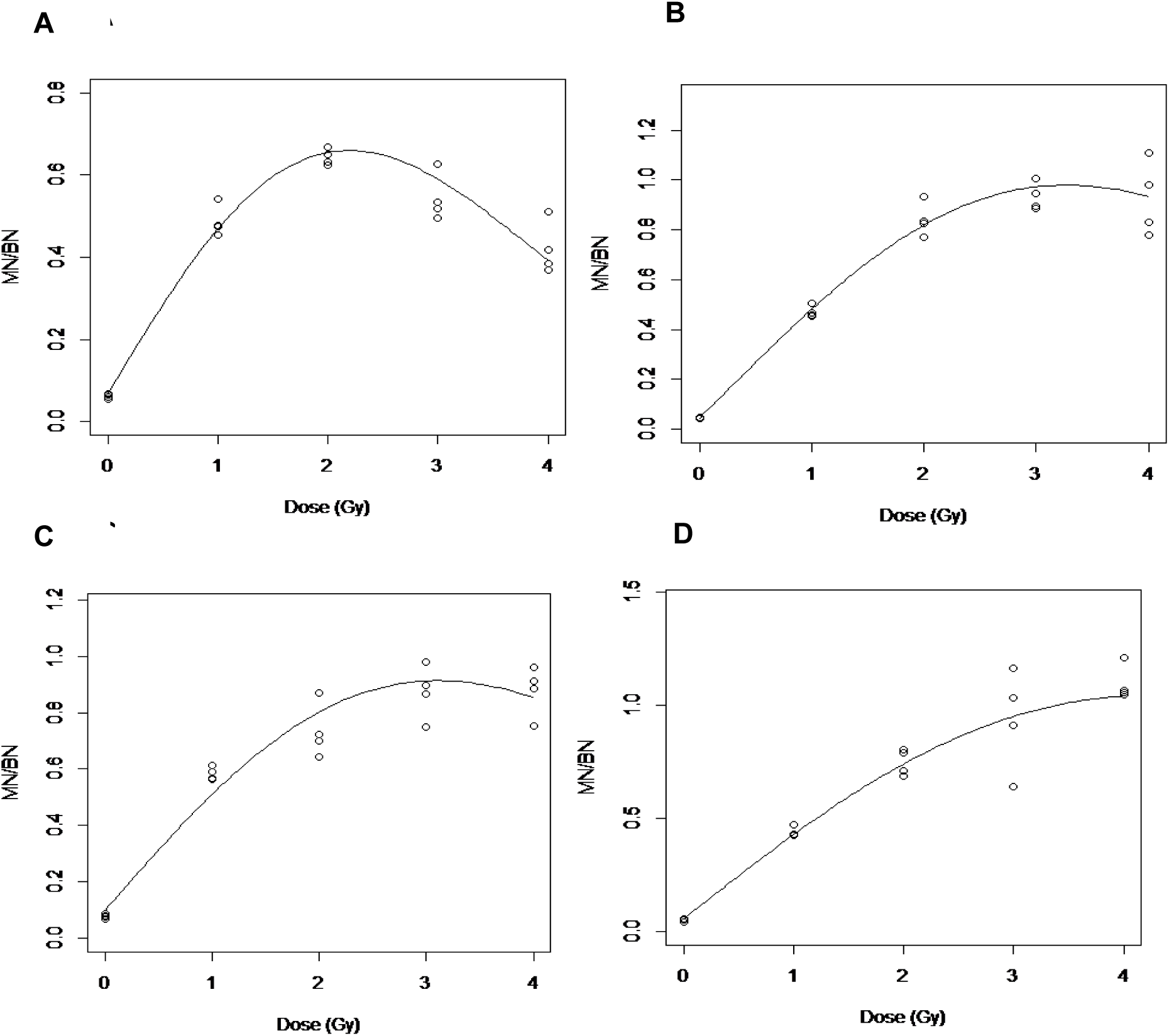
Dose effect curves after neutron irradiations. A. Standard CBMN assay; B. Caffeine CBMN assay; C. ZM447439 CBMN; D. Caffeine & ZM447439 CBMN assay.

**Table 1.**
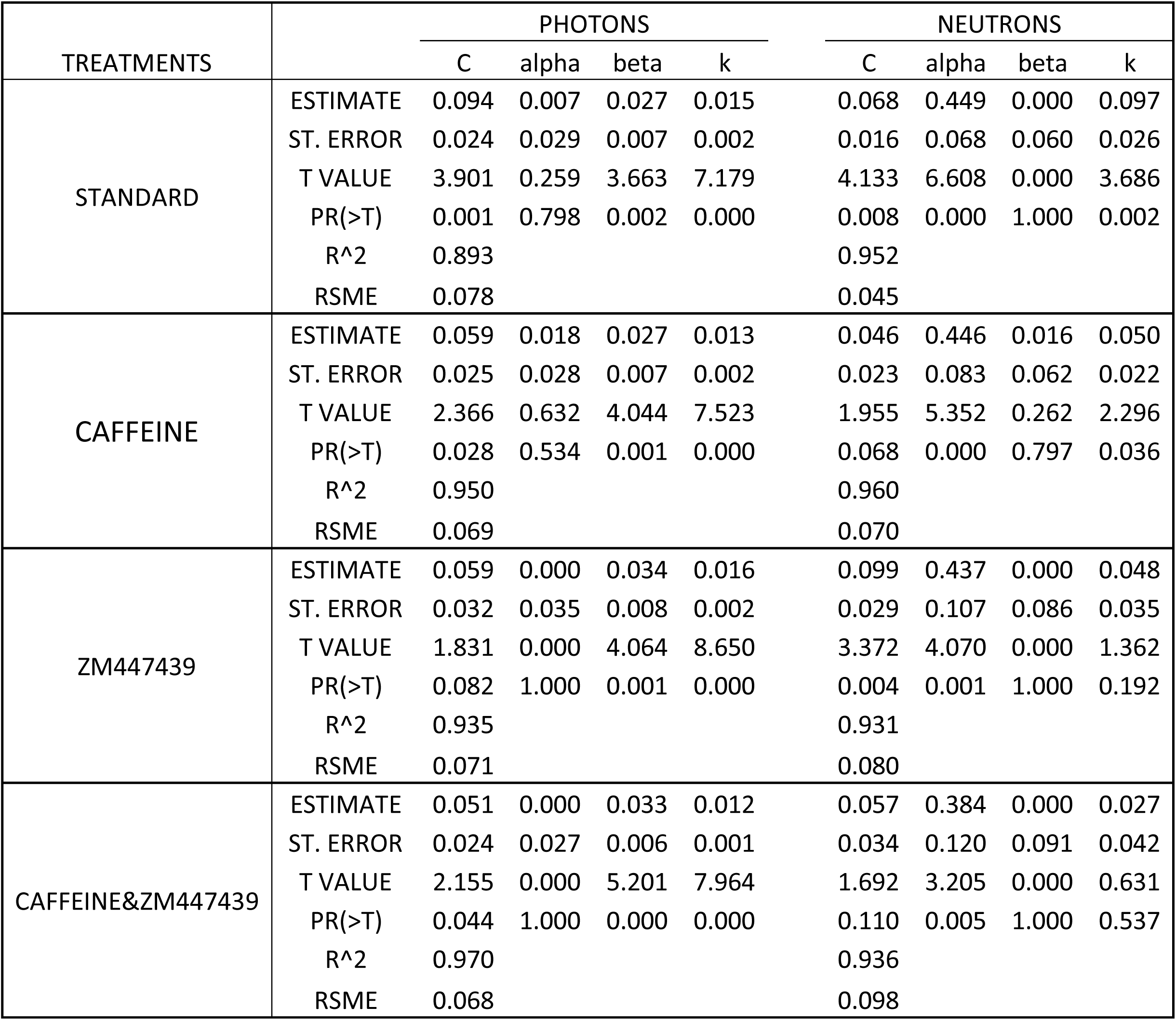

## Discussion

Dose assessment using cytogentic methods has limitations at high radiation doses because of the inhibitory effect on cell proliferation exerted by radiation (1). Therefore, most biodosimetry studies have not attempted to investigate doses higher than 5 Gy for low LET radiation, and even lower doses when high LET irradiations, such as with neutron exposures, are involved. In the present study, the goal was to modify the CBMN assay protocol to achieve a dose-effect curve that increased mononotonically for doses up to 10 Gy of low LET radiations and make the assay applicable to high LET radiations.

It has been described that extension of culture time can allow more cells to reach metaphase in the case of the dicentric assay (25)(26). Culture time extension has also been applied to the CBMN assay, where a clear increase in Mn/BN cells has been observed after neutron irradiations (27). Additionally, it has been shown that the combination of the culture time extension along with caffeine treatments increases the number of heavily damaged cells reaching metaphase (18)(19)(20). This study builds on these previous observations and shows that the combinations of caffeine/ ZM447439 along with an extended culture time can increase the dose detection limit thereby making the assay applicable to a wider range of radiation exposure scenarios.

As seen in Figure 1 at doses of 8 Gy and 10 Gy, a higher frequency of Mn/BN was observed in caffeine and ZM447439 treated cultures compared to the untreated ones. At doses lower than 4 Gy photons or 2 Gy neutrons, the linear increase of Mn frequency with the dose reflects the fact that the cells are still able to progress through the cell cycle. Here we show that inhibition of the G2/M arrest and spindle checkpoint allow cells heavily damaged by relatively higher doses of radiation to reach cytokinesis. As has been showed before in dicentric assay (18), a positive effect has been seen extending the caffeine treated cultures in the CBMN assay. A significant increase of the Mn/BN was found with 74 hours caffeine treated cultures when they were compared to the standard protocol, but no significant benefit has been observed when the 74 hrs was extended to 78 hrs. Although the Mn/BN increased when caffeine was added to the cultures, a saturation was observed at higher doses; for this reason, we proposed a double inhibition of the G2/M and spindle checkpoint to get an increasing ratios of Mn/BN as dose increases even at high doses of radiation.

In Figure 2, a comparison between the four different treatment protocols (Caffeine, ZM447439 alone or together) explored in this work can be seen at different doses after either photon or neutron irradiations. A similar dose response has been seen after both types of radiation by the different assay modifications. At lower doses, neither the single inhibition of the G2/M or the spindle checkpoints or the double inhibition by the combination of caffeine and ZM447439 produces a significant difference in Mn/BN frequencies. This lack of effect is likely because at lower doses the frequency of Mn is relatively low, and damaged cells are not negatively selected by the cell cycle checkpoints. At higher doses when caffeine was added to the cultures almost no turnover has been observed (Figure 2), although a saturation on the highest doses can be seen. For this reason, we expected that the combination of the caffeine and ZM447439 for G2/M and spindle checkpoint abrogation could allow micronucleated cells to reach kyriokinesis. When the results between the single inhibition checkpoints are compared we have seen that with the G2/M abrogation by caffeine, higher frequencies of Mn/BN cells as well as more binucleated cells could be scored (Figures 2 and 3). This is probably because the G2/M checkpoint occurs earlier in the cell cycle so the number of cells reaching this checkpoint is larger, meaning that the number of cells that reach the spindle checkpoint when the G2/M is active after high doses is reduced.

While the abrogation of the spindle checkpoint alone is not very useful after photon irradiaton, the double inhibition appears more effective that the single G2/M abrogation; his is likely because the pool of cells that reach metaphase is significantly higher when the G2/M checkpoint is also inhibited.

When the curve fitting parameters (Eq. 1) are compared between different treatments, it can be seen that the lower k values are achieved with double checkpoint inhibition treatments for both photons and neutrons (Table 1).

At the highest doses assessed in the present study, 10 Gy in the case of photons and 4 Gy of neutrons, when double inhibition is compared to the standard protocol the observed frequency of Mn/BN is 2- and 2.5-fold lower with the standard protocol, respectively. In contrast, the standard assay treatment shows a clear turnover of Mn/BN yields to high doses after exposures to both types of radiation (Fig. 2, 3 and 4) due to the cell cycle arrest.

As has been widely observed, low-LET radiation dose responses are better fit using a linear quadratic model, rather than by a linear one. In the data presented here for photons, in some cases the best-fit linear coefficient, alpha, was equal to zero. This is probably because few values for the lower doses have been assessed, because the main goal of this study was to optimize the assay for high doses.

For the neutron exposures, in most of the dose response curves, the best-fit quadratic coefficient (beta) was 0, indicating that the dose response is consistent with linearity.

In the present study, after neutron exposures, huge differences in Mn/BN have been seen with single abrogation checkpoints, with even larger differences when double abrogation was induced. This is probably related to the cell cycle arrest produced by the complex aberrations and non-random distribution of energy from high-LET radiation exposure, which caused increased perturbation in the progression of the cell-cycle (28)(29)(30). For this reason, most of the studies performed with neutron irradiations rarely use doses higher than 2 Gy (12)(22)(30).

For high LET dose estimation, several studies suggested the use of chemically induced premature chromosome condensation as a method to score aberrations in G2 phase (29)(31). But another way to analyze the retained cells in G2 is the abrogation of the checkpoints. In this study we demonstrated that by this checkpoint abrogation approach reliable dose assessment could be performed following exposure with up to 4 Gy of neutrons and 10 Gy of photons. Likewise, using the modifications of the CBMN assay presented here, the assay shows promise as a methodology that could be used in high throughput scenarios.

## Conclusion

In the present work, we have optimized the standard CBMN assay for high-dose photon and neutron exposures through the addition of caffeine and ZM447439 (an Aurora kinase inhibitor) to the blood cultures at selected times. This approach increases the resolution resolution of the CBMN assay at higher doses, specifically up to 10 Gy and 4 Gy from photons or neutrons, respectively. This implies that the CBMN assay can be reliably employed to assess the absorbed dose even in realistic exposure scenarios in which individuals are likely to be exposed to relatively high doses of low and/or high LET radiation.

## Author Contributions

MPC, JRP, LC, IS, AH, GG, and DJB conceived the study and prepared the manuscript. MPC, JRP, LC, AH, and GG performed the experimental studies. IS carried out the data analysis. All authors contributed to editing the manuscript.

## Acknowledgments

This work was funded by a pilot grant from the Opportunity Funds Management Core of the Centers for Medical Countermeasures against Radiation, National Institute of Allergy and Infectious Diseases; grant number U19AI067773. The funders had no role in study design, data collection and analysis, decision to publish or preparation of the manuscript.

The authors would like to acknowledge Dr. Helen Turner, Dr. David Welch and Dr. Manuela Buonanno for their comments that greatly improved the manuscript

